# THiRST: Computer Vision Empowered Three-Dimensional High-Resolution Spatial Transcriptomics Analytics

**DOI:** 10.64898/2026.10.29.762926

**Authors:** Junchao Zhu, Yu Wang, Zhengyi Lu, Chongyu Qu, Yanfan Zhu, Yuechen Yang, Shilin Zhao, Haichun Yang, Yuankai Huo, Ruining Deng

## Abstract

High-resolution spatial transcriptomics (ST) preserves gene expression within tissue architecture but remains limited to individual two-dimensional (2D) sections, restricting analysis of continuous three-dimensional (3D) structures and cross-section consistency. We present THiRST, a computational framework for reconstructing multi-resolution 3D spatial transcriptomic representations from serial sections. THiRST combines coarse-to-fine morphology-guided registration, spatial matching of ST units, cross-section association of segmented structures, and integration of Visium V2 and high-definition ST. In serial kidney sections, patch-level refinement increased the same-cluster proportions of matched units at adjacent offsets from 63.8% and 65.6% to 70.7% and 70.0%, respectively, while improving expression concordance. The reconstructed correspondences enabled 3D glomerular and object-level expression analysis, with 54.1% of differential genes detected only in 3D. Cross-platform integration further aligned V2 and high-definition measurements in a shared expression space. THiRST demonstrates the feasibility of object-level 3D ST analysis from serial 2D sections and underscores the importance of accurate local registration.

## 1. INTRODUCTION

Spatial transcriptomics (ST) measures gene expression while preserving tissue spatial organization, directly linking molecular signals to histological structures [1]. High-resolution platforms such as Visium HD further reduce measurement units from 55 *µ*m spots to 2 *µ*m bins, approaching the single-cell scale [2]. However, each section remains two-dimensional, whereas renal structures such as glomeruli, tubules, and blood vessels extend continuously in three dimensions (3D) and may exhibit depth-dependent pathological changes [3, 4]. Serial-section registration therefore provides a practical route to three-dimensional ST reconstruction [5]. Prior studies have reconstructed virtual tissue volumes [6], predicted expression in unmeasured sections [7, 8], or jointly modeled multiple sections using deep learning [9, 10]. Yet the reproducibility of high-resolution ST across adjacent sections remains unclear, as sparse transcript counts, tissue deformation, and batch effects can introduce variation even at corresponding anatomical locations. Reliable 3D reconstruction therefore requires precise spotor bin-level correspondence across sections, together with harmonization across platforms of different resolutions to ensure spatial consistency.

To address these challenges, we propose THiRST, a computer vision-driven framework for 3D ST analysis from serial tissue sections (Figure 1). Serial kidney sections profiled with Visium V2 and Visium HD are aligned using morphologyguided coarse-to-fine registration [11], followed by spatial matching of spots and bins across tissue sections. Renal structures are segmented and linked across sections to reconstruct 3D objects [12], while CCA integrates V2 and HD data into a shared expression space [13, 14]. We assess cross-section reproducibility through cluster agreement and expression correlation between matched locations, showing greater consistency between adjacent sections and after local registration refinement. We further reconstruct glomeruli in 3D and aggregate expression at the object level. Compared with singlesection analysis, 3D analysis identifies more differentially expressed genes while preserving coherent spatial marker patterns, highlighting the value of volumetric ST analysis. Our contributions can be summarized as threefold:

- We propose THiRST, a 3D ST analysis framework from conventional serial tissue sections, integrating morphologyguided coarse-to-fine registration, structural segmentation, 3D object association, and Visium V2/HD integration within a unified coordinate system.
- We quantitatively evaluate high-resolution ST reproducibility across serial sections and between high- and low-resolution platforms using spot- and bin-level correspondence, and examine the impact of registration accuracy on these measurements.
- We demonstrate the feasibility of 3D object-level ST analysis using glomeruli, showing volumetric analysis identifies more disease-associated differentially expressed genes than conventional single-section 2D analysis.

**Fig. 1.**
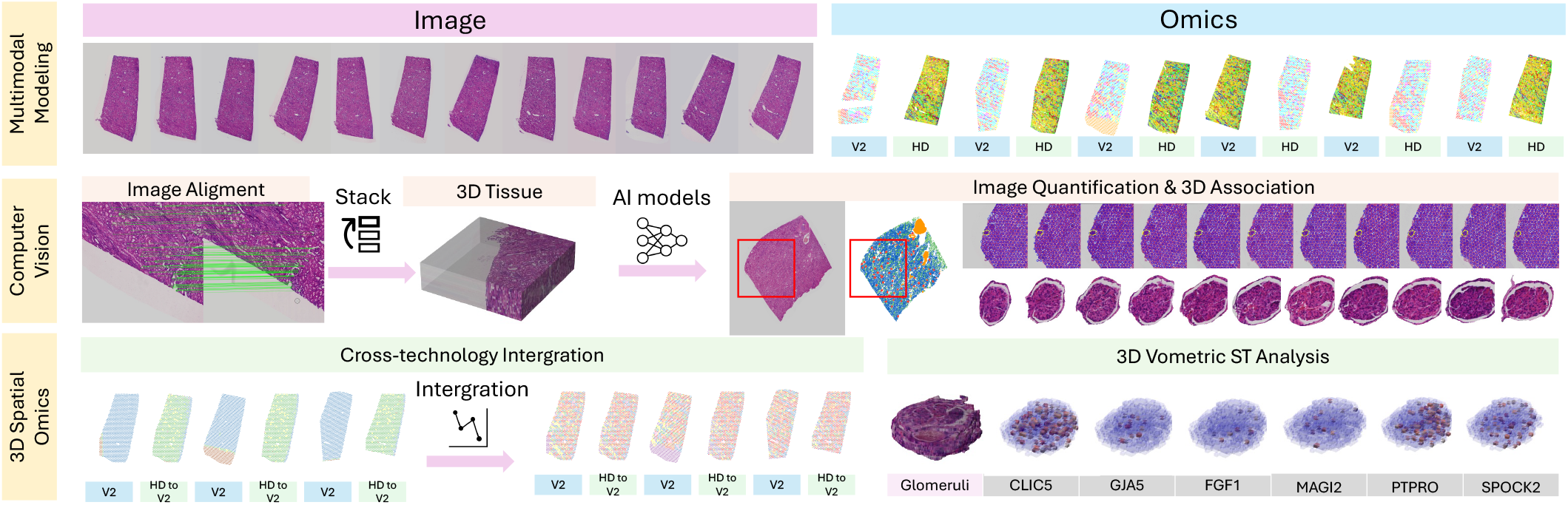
Overall pipeline of THiRST. THiRST integrates serial histology and multi-platform ST data into a unified 3D representation. V2 spots and HD bins are mapped into a 3D coordinate system to reconstruct volumetric expression patterns. CCA harmonizes low- and high-resolution ST data into a shared transcriptional space for tissue- and object-level analysis.

## 2. METHOD

### 2.1. Coarse-to-Fine 3D Registration and Harmonization

The input consisted of an ordered stack of *N* consecutive tissue sections, 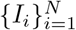, each associated with a whole-slide image (WSI), ST measurements, and a gene-expression matrix. Visium V2 locations were represented as circular spots, whereas high-resolution ST units were modeled as rectangular bins. The middle section, *I_r_* with *r* = ⌊*N/*2⌋, was selected as the reference plane. All WSIs, ST contours, and morphological annotations were mapped to this common coordinate system using a coarse-to-fine registration strategy. At the global stage, adjacent WSIs were aligned using feature-based affine registration followed by multi-resolution refinement. Let **A**_*b*←*a*_ ∈ ℝ^3*×*3^ denote the homogeneous affine transformation from section *I_a_* to adjacent section *I_b_*. The global mapping from *I_i_* to the reference plane was obtained by composing pairwise transformations along the stack:

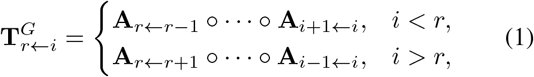

where reverse mappings were obtained by inverting the corresponding affine matrices. Since global alignment may leave residual deformation around fine structures, a second patch-level refinement was applied. Patch centers {**c***_p_*} were sampled from ST barcode locations, and corresponding local fields of view were extracted across the registered stack. Each local stack was initialized by feature matching and refined using non-rigid ANTs registration, yielding a local deformation mapping **Φ***_i,p_*. Each point **x** on section *I_i_* was assigned to the nearest patch center in the globally aligned space:

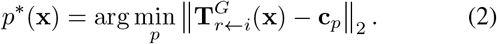

The final 3D coordinate was defined as

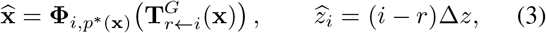

where Δ*z* denotes the physical spacing between adjacent sections. RGB images were resampled using bilinear interpolation and segmentation masks using nearest-neighbor interpolation. ST units were represented by polygonal contours, and the composite transformation was applied to all contour vertices to preserve spatial correspondence among histology, morphology, and transcriptomic measurements in 3D space.

### 2.2 Cross-Section Reproducibility of HD ST

Following spatial harmonization, the *m*-th ST measurement unit on section *I_i_* was represented by a polygonal footprint *P_i,m_* in the reference plane. Spatial correspondence between two units was quantified by intersection over union (IoU), defined as 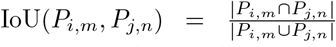 To reduce the cost of large-scale contour matching, candidate pairs that could not overlap were discarded using bounding-box intersection and centroid-distance tests. For each pair of sections, one-to-one correspondences were then obtained from the IoU matrix using the Hungarian algorithm, with*π*^*^ = arg max*_π_* ∑*_m_* IoU (*P_i,m_, P*_*j,π*(*m*)_). Assignments with IoU *< τ* were rejected, with *τ* = 0.01 used throughout. The same procedure was applied to adjacent and nonadjacent section pairs to assess reproducibility across section offsets. For cross-resolution comparison, HD bins were aggregated onto the V2 spot grid based on geometric overlap. Given the expression vector **x***_i,b_* of HD bin *b*, its overlap weight and contribution to V2 spot *s* were defined as

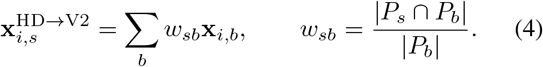

All datasets were restricted to a shared gene set and processed using the same normalization, feature-selection, principal-component analysis, and joint clustering pipeline. Reproducibility was assessed at two complementary levels. The matched-pair set for section offset *d* = |*i* − *j*| was denoted by ℳ*_d_*, and *y_i,m_* represented the joint cluster assignment of unit (*i, m*). Cluster concordance was quantified as

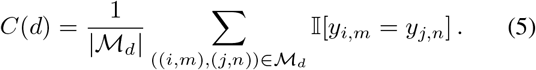

Expression-level reproducibility was evaluated using the normalized expression vectors 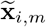 over the shared gene set. Pearson correlation was computed for each matched pair and summarized within each section offset:

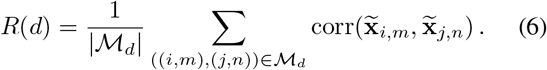

Here, *C*(*d*) measures agreement in discrete transcriptomic identity, whereas *R*(*d*) captures continuous similarity of gene-expression profiles. Both metrics were computed separately after global registration and patch-level refinement and were stratified by section offset *d* to quantify the effect of local alignment on cross-section reproducibility.

### 2.3 3D Morphology–Transcriptomics Analysis

To construct the 3D morphological layer, each WSI was tiled using a tissue foreground mask and processed with the scale-aware Omni-Seg model. Six renal structures were segmented: distal tubules, proximal tubules, glomerular capsules, glomerular tufts, vessels, and peritubular capillaries. Tile-level predictions were fused at the WSI level, and the resulting contours were mapped to the common 3D coordinate system using the composite transformations. Objects of the same class on adjacent sections were linked across depth when their contours satisfied IoU *>* 0.01, forming 3D tracks. Morphology–transcriptomics concordance was then quantified using volumetric IoU. Let *M_i,t_* denote the spatial support of morphological class *t* on section *i*, and *Q_i,c_* the domain of transcriptomic cluster *c*. Their 3D overlap was computed as 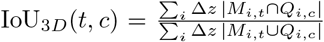. For reconstructed structures such as glomeruli, was aggregated across ST expression units overlapping each 3D object. Let Ω*_k_* denote the overlapping units for object *k*. Each unit *u* was weighted by the fraction of its footprint covered by the object mask, yielding the object-level expression vector:

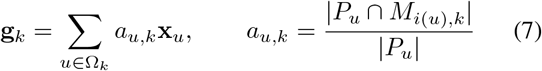

The object-level profiles were normalized for comparison between diabetic nephropathy and healthy tissue, with sectionwise analysis used as the 2D reference. For joint Visium V2 and HD-ST analysis, shared highly variable genes were integrated using canonical correlation analysis (CCA), followed by anchor-based clustering in the shared latent space. Cluster labels and expression patterns were then mapped back to the unified 3D coordinates. Morphology–transcriptomics concordance, glomerular expression patterns, and 2D versus 3D differential-expression analysis were used to evaluate the feasibility of serial-section 3D ST reconstruction.

## 3. RESULTS

### 3.1 Refined Cross-Section ST Concordance

THiRST established spatial correspondences between ST units in the unified 3D space, enabling cross-section reproducibility to be assessed at the cluster and gene-expression levels (Fig. 2). With global registration alone, 63.8% and 65.6% of matched units at the two adjacent offsets belonged to the same transcriptomic cluster. Patch-level refinement increased these proportions to 70.7% and 70.0%, respectively (Fig. 2a–b). Improvements were observed across most offsets and were strongest between nearby sections, while concordance generally decreased with increasing section separation. Expression-level concordance showed a similar trend (Fig. 2c–d). Patch-level refinement shifted correlation distributions toward higher values, whereas more distant pairs showed lower and broader correlations. The uneven improvement across offsets suggests that refinement mainly corrected residual spatial misalignment rather than biological heterogeneity or sparse transcript capture. Overall, local refinement improved cross-section ST concordance and provided more reliable correspondences for 3D analysis.

**Fig. 2.**
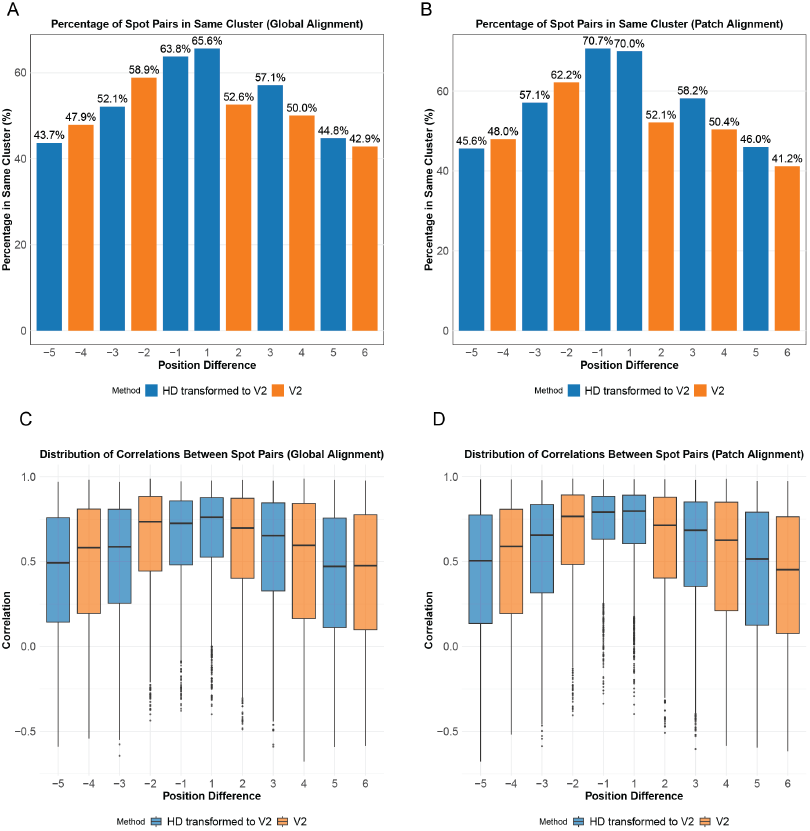
Patch-level refinement improves cross-section ST concordance. **(a–b)** Same-cluster proportions after global registration and patch-level refinement. **(c–d)** Geneexpression correlations under the same settings, with the largest improvements in nearby sections.

### 3.2 3D ST Analysis from Serial Sections

Following spatial harmonization, glomerular contours and their associated HD-ST signals were linked across consecutive sections to form 3D objects (Fig. 3a). The reconstruction retained the morphological continuity of each glomerulus across depth and enabled the spatial distributions of representative genes, including CLIC5, GJA5, FGF1, MAGI2, PTPRO, and SPOCK2, to be visualized within the same 3D structure. This result demonstrates that morphology and gene expression acquired from conventional 2D sections can be represented jointly at the 3D object level. Differentialexpression results between diabetic nephropathy and healthy tissue were compared at the glomerular, podocyte, and glomerular endothelial cell levels (Fig. 3b). Under the current settings, 54.1% of differential genes were detected only by 3D analysis, 36.9% by both 2D and 3D analyses, and 9.0% only by 2D analysis. By aggregating measurements across multiple depths of the same reconstructed structure, 3D analysis reduced dependence on a single sectioning plane. These results support the analytical feasibility of object-level 3D ST analysis, rather than implying that all additional 3D-specific genes are necessarily more biologically accurate.

**Fig. 3.**
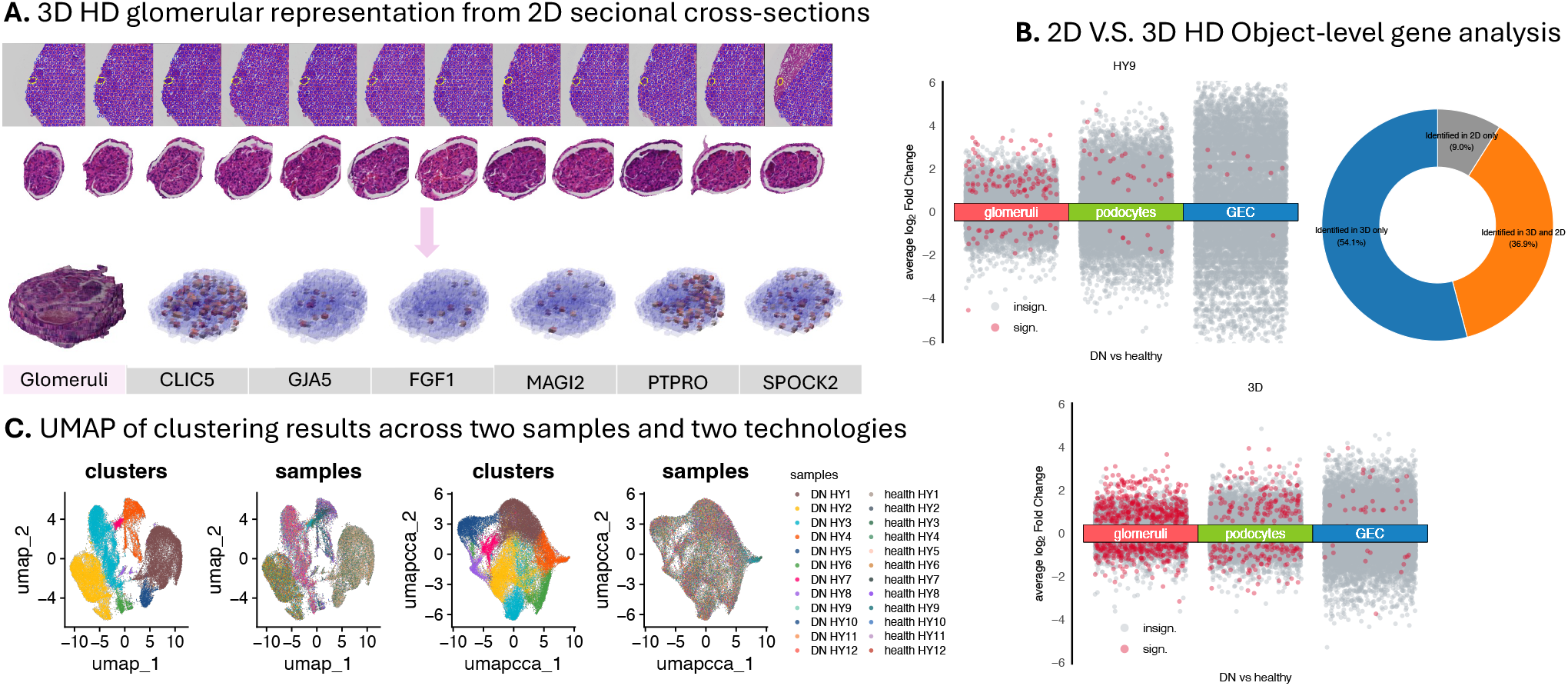
Object-level 3D analysis of serial ST sections. **(a)** 3D reconstruction of glomerule and representative HD-ST geneexpression patterns. **(b)** Differential genes identified by 2D section and 3D object-level analyses of diabetic nephropathy and healthy tissue. **(c)** UMAP embeddings of Visium V2 and HD-ST measurements before and after CCA integration.

Cross-platform integration assessed whether Visium V2 and HD-ST could share a common expression space (Fig. 3c). Before integration, UMAP showed clear section- and technology-associated separation. After CCA, measurements from different sections and platforms were intermingled while preserving distinct transcriptomic clusters. This result supports the compatibility of the two spatial resolutions within a shared transcriptional representation. Together with 3D reconstruction and volumetric expression aggregation, these findings demonstrate the feasibility of morphology– transcriptomics analysis from serial 2D ST sections.

## 4. CONCLUSION

We present THiRST, a unified computational framework that integrates morphology, transcriptomics, and spatial geometry within a common 3D coordinate system to construct multiresolution tissue representations from conventional serial 2D sections. By combining coarse-to-fine morphology-guided registration, cross-section structural association, and crossplatform harmonization, THiRST supports tissue-scale reconstruction and object-level molecular analysis in a single workflow. Analyses of kidney tissues demonstrate its ability to improve spatial correspondence across sections and enable 3D structural reconstruction, object-level expression comparisons, and joint representation of different ST platforms. Together, THiRST provides a practical and extensible foundation for transforming existing 2D ST data into volumetric views of tissue structure and molecular organization, with the potential to advance 3D biomarker discovery.

## 5. COMPLIANCE WITH ETHICAL STANDARDS

The private kidney dataset used in this study was collected under approval from the Institutional Review Board (IRB) of Vanderbilt University Medical Center (IRB No. 230223).

## 6. ACKNOWLEDGMENTS

This research was supported by the National Institutes of Health (NIH) under grant R01DK135597, with $20000, representing 66% of the total project costs. The content is solely the responsibility of the authors and does not necessarily represent the official views of the NIH. This material is based upon work supported by the U.S. National Science Foundation under Award Number 2434229. A total of $5000 representing 17% of the total project costs, was financed with this funding. Any opinions, findings, and conclusions or recommendations expressed in this material are those of the author(s) and do not necessarily reflect the views of the U.S. National Science Foundation. A total of $5000 representing 17% of the total project costs, was financed with non-governmental funds.

